# Staring at the naked Goddess. Unraveling structure and reactivity of Artemis endonuclease interacting with a DNA double strand

**DOI:** 10.1101/2021.06.02.446693

**Authors:** Cécilia Hognon, Antonio Monari

## Abstract

Artemis is an endonuclease responsible for breaking hairpin DNA strand during immune system adaptation and maturation as well as the processing of potentially toxic DNA lesions. Thus, Artemis may be an important target in the development of anticancer therapy, both for the sensitization of radiotherapy and for immunotherapy. Despite its importance its structure has been resolved only recently, and important questions concerning the arrangement of its active center, the interaction with the DNA substrate, or the catalytic mechanism remain unanswered. In this contribution, by performing extensive molecular dynamic simulation, both classically and at hybrid quantum mechanics/ molecular mechanics level, we evidence the stable interaction modes of Artemis with a model DNA strand. We also analyze the catalytic cycle providing the free energy profile and key transition states for the DNA cleavage reaction.

## 1. Introduction

Artemis is an endonuclease [1–3] which plays a fundamental role in processing DNA strands in immune system cells allowing their recombination and hence the maturation of the immune response. Indeed, Artemis is present in B and T cell and its catalytic activity concerns the opening of the DNA hairpins in the V(D)J process. V(D)J [4,5] is the mechanism through which immune system cells assemble variable (V), diversity (D), and joining (J) gene sequences responsible of the production of Immunoglobulins (Igs) and T-cell receptors that can recognize a large number of antigens. Thus, Artemis is a key component allowing the somatic development of Igs and T cells in superior animals. Its activity is also finely regulated by specific cellular pathways, and namely Artemis forms a complex with the DNA dependent protein kinase catalytic subunit (DNA-Pkcs), which induces phosphorylation and activation of the enzyme, most probably by exposing its catalytic site [6,7]. Thence the activation of Artemis allows the opening of the DNA hairpins that have been generated by the recombination activated gene during V(D)J [8]. Importantly, since the Artemis/DNA-Pkcs is the only protein ensemble able to effectively open DNA hairpins [9], functional inhibiting mutations in either partner effectively block B and T cell maturation and leads to phenotypes showing severe combined immunodeficiency [10], and to the accumulation of hairpin coding ends in thymocytes [9]. In addition to its fundamental role in immune system maturation, Artemis has also been associated to the capacity of processing DNA lesions, and in particular strand breaks, hence participating to the maintaining of genome stability [11–13]. Indeed, non-functional mutations of Artemis have been correlated to an increased radiosensitivity [10] of B and T cells, suggesting its participation to the strand break repair machinery [14]. Notably, it has been shown in cellular models that Artemis is recruited at the site of DNA damages and acts in a way that is similar to non-homologous end joining pathways to assure repair [15,16]. Interestingly, the interplay between DNA repair and immune system maturation has also been highlighted, in the sense that DNA repair machinery can participate to the last steps of the V(D)J process [10].

Artemis shows sequence similarity and homology with a number of RNA ribonuclease, and two human nucleases of the SNM1 family: SNM1A and SNM1B/Apollo, hence it belongs to the metallo β-lactamase class [6]. Despite obviously similarities Apollo, the twin counterpart of Artemis, presents only an exonuclease activity and the density of surface positive charges, allowing for the efficient binding of the DNA substrate, is strongly reduced [17].

As a matter of fact, the function and the biological role of Artemis could potentially result in a most ideal target candidate for chemotherapy or radiotherapy sensitization [18,19]. Furthermore, its participation in the immune system maturation could also point to the possibility of exploiting its action in immunotherapy strategies. However, the structure of the catalytic and DNA binding domains of Artemis has been resolved only very recently. Furthermore, while the protein was correctly resolved no bound nucleic acid was shown in the obtained crystallographic structure and some key partners in the catalytic site were not resolved [20]. The lack of a precise structural and dynamic characterization of Artemis behavior clearly hampers the rational development of potential inhibitors that could be used as chemotherapeutics or as radiotherapy sensitizers.

From a biophysical point of view, different structural characteristics of Artemis have been pinpointed, notably by Karim *et al*. [20], these include the presence of a zinc finger and a surface groove presenting a high density of positively charged aminoacids, mainly lysines, which should be essential for DNA recognition and stabilization. From a chemical point of view, exo- or endonuclease activity is usually performed by a catalytic site featuring a metal or a metal cluster, usually Zn^2+^ or Mg^2+^, which acts through mono or bimetallic pathways [21]. In the case of Artemis contrasting structural evidences have been reported, and in many cases only one Zn^2+^ ion has been located in the active site. However, also considering the similarity with the other members of the SNM1 family, exerting bimetallic activity, it seems reasonable that a bimetallic cluster should be present in the biological active form. Furthermore, despite the presence of zinc in the crystal structure, the evidence of sustained activity of the enzyme in a magnesium buffer [20], strongly points towards the possibility of a bimetallic (Mg^2+^)_2_ active center.

In the present contribution we resort to high-level molecular modeling and simulation to explore the inherent structural properties of Artemis and its interaction with DNA double strand. In addition, we will also examine the reaction profile leading to the catalytic DNA strand break confirming the activity of the (Mg^2+^)_2_ cluster. To this aim, we consider a multiscale approach combining both full-atom classical molecular dynamics (MD) simulations exceeding the μs timescale to explore the DNA binding and its structural evolution, with hybrid quantum mechanics/ molecular mechanics (QM/MM) approach. The latter involves enhanced sampling strategies, namely umbrella sampling (US), to obtain the free energy profile (FEP) of the most relevant and critical steps of the DNA cleavage. Hence, through our work we provide a unified vision of Artemis functioning, resolving the still elusive or contrasting hypotheses on this crucial protein.

## 2. Results

We firstly performed MD simulation of the unbound wild type form of the catalytic domain of human Artemis starting from the structure reported by Karim *et al*. (pdb: 6wo0) [20]. In order to better describe the bimetallic cluster constituting the catalytic site we included a second Mg^2+^ ion together with bonding constraints on the two metals to avoid instability due to positive charge repulsion. As shown in SI, and as witnessed by the small value of the root mean square deviation (RMSD) all along the trajectory the protein structure is stable with deviations plateauing at 3.8 ± 0.6 Å. Most importantly, the main secondary and tertiary structural motifs, in particular two extended β-sheet regions connected by loops and α-helices, are preserved all along the dynamic. A positively charged groove, composed mainly of lysine and approaching the catalytic active site is also evident and solvent exposed constituting an optimal target for the DNA binding. Interestingly, along the MD simulation of the isolated protein the charged groove also shows a remarkable stability with only limited oscillation of its width and depth, an occurrence that could be ideal to allow an efficient binding with the nucleic acid.

In the crystal structure of pdb: 6wo0 the bound DNA strand was not resolved. Hence, we firstly proceeded to a protein/DNA docking analysis to identify suitable starting poses. As shown in Figure 1A, and unsurprisingly, the most favorable pose places the DNA backbone in the positively charged groove of the protein. Although, electrostatic interactions appear evident and favorably to drive the binding, it is also clear that the ideal straight arrangement of the DNA strand used for the docking is not able to maximize the contacts and lead to a compact complex. These general observations are indeed confirmed by the analysis of the MD simulations performed from the docking pose, which however points to the establishment of a persistent complex remaining stable for the entire simulation reaching 2 μs. Of note, an independent replica, also spanning 2 μs, has been performed giving similar results (see SI). As shown in Figure 1B, the time evolution of the RMSD for the global Artemis/DNA complex remains globally moderate and is stabilizing at around 6.5 ± 0.9 Å. On the contrary, the contribution due to the nucleic acid is clearly more important, and while averaging at 11.9 ± 2.0 Å, it also shows more pronounced oscillations, which are indicative of a greater flexibility and the possible coexistence of slightly different arrangements.

**Figure 1.**
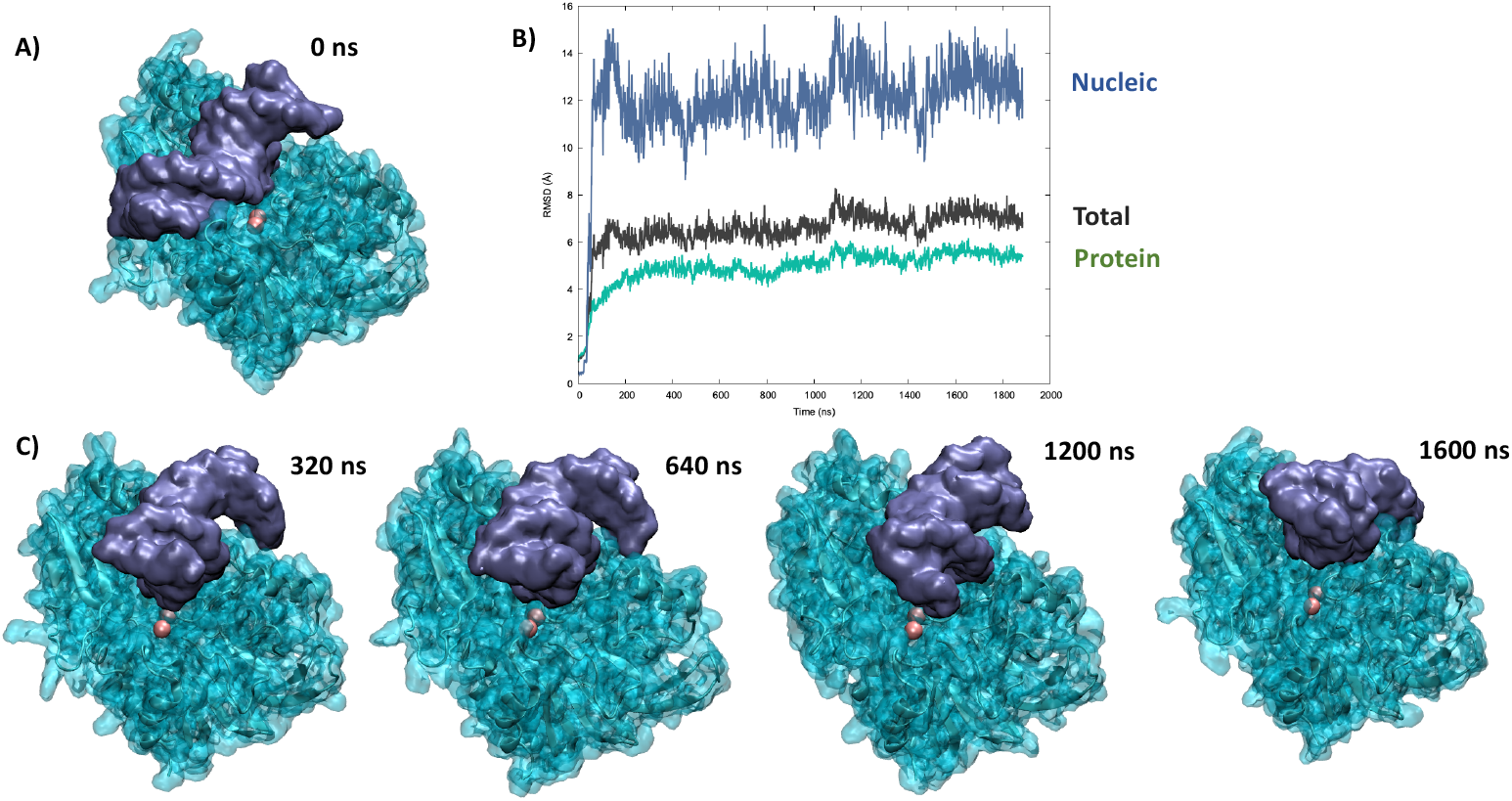
A) Most favorable pose resulting from the docking of the crystal structure of Artemis with a DNA double strand, B) Time evolution of the RMSD for the MD simulation of the Artemis/DNA complex, and C) Representative snapshots extracted at different time frames of the MD simulation.

A part from the analysis of the RMSD, the reorganization of the bound DNA strand is also clearly evidenced by the visual analysis of the trajectory, as shown by the snapshots reported in Figure 1C. Indeed, it is evident that the strand tends to bend considerably to accommodate in the positively charged groove and maximize the contact. This fact is also evidenced by the analysis of the DNA structural parameters, and notably the global bending, performed with Curves+ and reported in the SI. Interestingly, the bound nucleic acid also tends to show sliding and rotation in the groove, which while maintain a stable complex may also allow a certain flexibility and the possibility to present different phosphate units to the vicinity of the catalytic site. This aspect can also be related to the biological role of Artemis, which requires the ability to cleave unspecific DNA sequences either for repair of V(D)J maturation.

Looking more in detail to the specific interactions stabilizing the DNA/Artemis complex (Figure 2) one may see that the binding is mainly driven by extended salt bridge involving the DNA backbone phosphate and the lysine populating the positively charged groove. This can be clearly appreciated by examining the radial distribution function of the distance between the negatively charged oxygens of the phosphate groups and the hydrogen atom of the positive ammonium moiety in Lys. As shown in Figure 2, the distribution presents a very intense and sharp peak at 2 Å that is indicative of the formation of rather rigid and persistent interactions. Note that a secondary peak is also observed at larger distance which can be safely attributed to the other ammonium hydrogens. In the snapshot reported in Figure 2, one can also clearly identify that the Lys later chains are indeed interacting strongly with both the DNA major and minor groove hence providing strong anchoring points. Interestingly, the global architecture, involving the penetration of Lys in the grooves, is also compatible with a global combined rotation and translation of the nucleic acid that is reminiscent of a “corkscrew” translation. This movement was observed in one of the replica MD simulations, see the video in Supplementary Information, and clearly allows to expose different phosphate and glycosidic bonds to the vicinity of the active site while preserving the global stability of the complex and the favorable interactions. Once again, this observation is compatible with the recognized capacity of Artemis to process different DNA sequences, notably to allow for the V(D)J maturation as well as to its recognized exonuclease activity that adds up to the more conventional endonuclease capacity [22].

**Figure 2.**
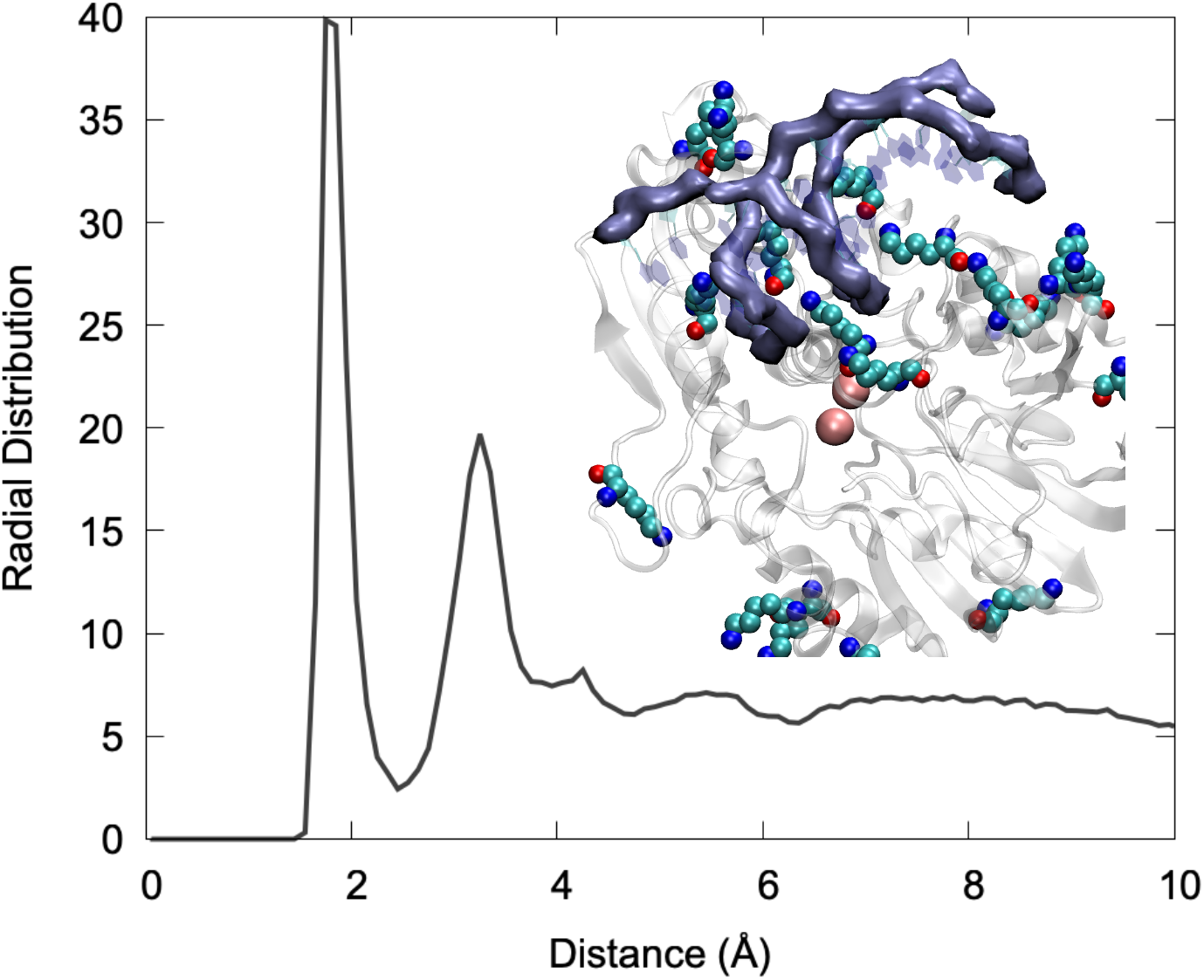
Radial distribution function between the negatively O1P and O2P atoms of the DNA back-bone and the HZ hydrogen of LYS ammonium group. A snapshot showing the interaction of LYS charged moieties (in van der Waals representation) with DNA backbone (highlighted with the purple surface) is also shown in the inlay.

In Figure 3, we may evidence the main characteristic of the active site of Artemis. Notably, and in addition to the two Mg^2+^ ions, we can also identify negative, or electron rich ligands, that are necessary to stabilize the metal cluster. In particular a high density of histidine is present that are susceptible to interact with the magnesium cation. More importantly, the presence of two aspartates, ASP 61 and ASP 160, seam crucial, since the harder oxygen may be a better ligand for magnesium than nitrogen. The analysis of the active site, and in particular the lack of an additional Asp ligand compared to other endonuclease such as SNM1B/Apollo, also justify the more labile nature of the second Mg^2+^, which makes hard to resolve its position by crystallography. Importantly, the active site is also relatively solvent exposed and as a matter of fact water molecules interact strongly with the (Mg^2+^)_2_ cluster, as seen by the sharp peak present in the radial distribution function at 2 Å (Figure 3) and by the fact that in average 5 water molecules are in a radius of 3 Å from Mg^2+^, as can be inferred from the integral of the radial distribution function. In the same context, it is worth mentioning that while using classical MD we were only exploring pre reactive conformation, in which the phosphate group still lays relative further from (Mg^2+^)_2_, we may observe, as seen in Figure 3, an important stabilization of its distance. Note that in some instances of the second replica a switch of the closest phosphate oxygen has been observed, coherently with the global motion of the DNA strand described previously. Furthermore, Asp61 and His62, may also have a more stringent function than the simple stabilization of the bimetallic cluster and may instead play an active role in the activation of the water molecule attacking the phosphate bond by acting as hydrogen acceptor. This is also confirmed by the fact that the mutation of Asp 61 totally suppresses the catalytic activity.

**Figure 3.**
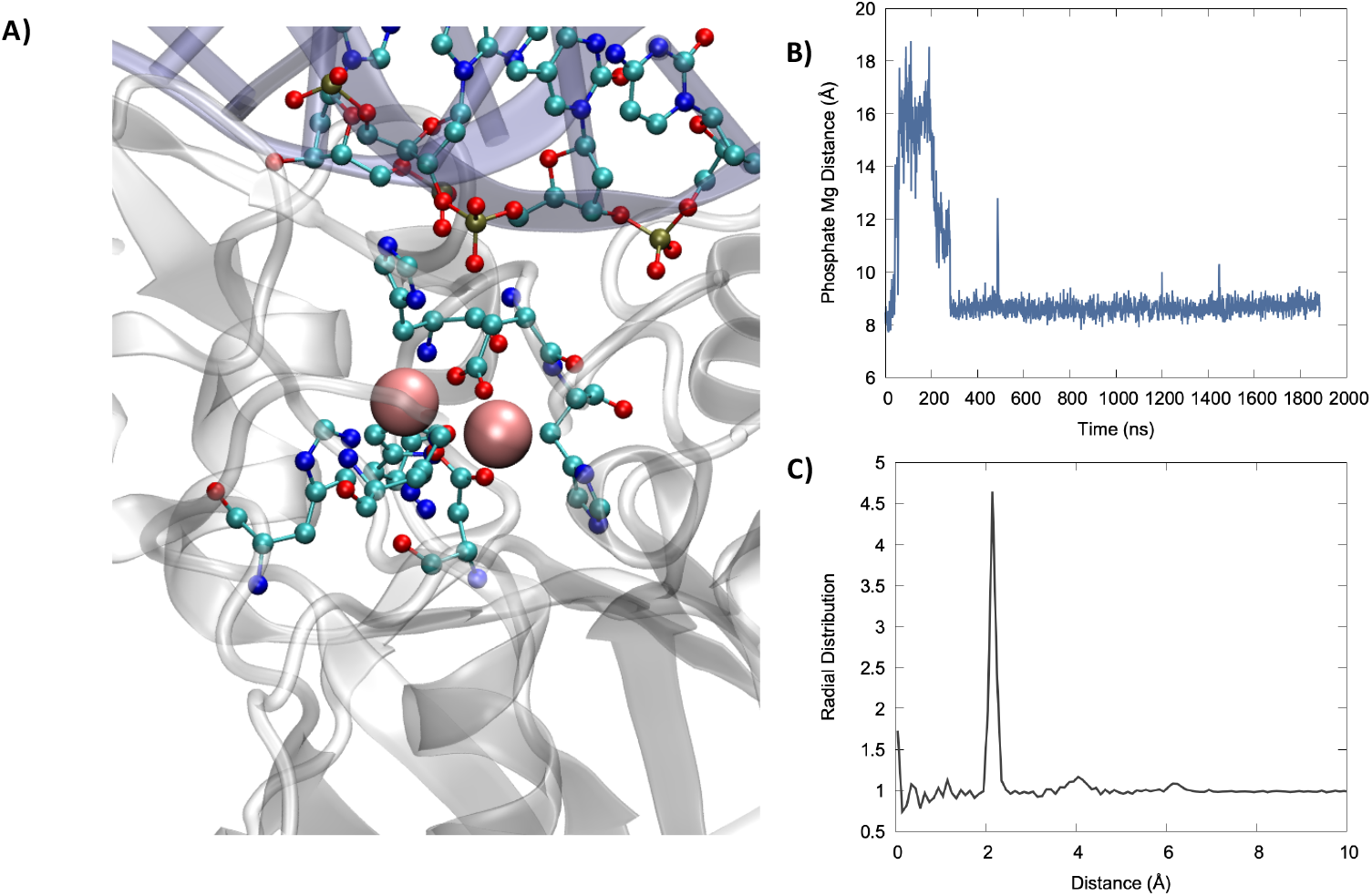
A) Representative snapshot highlighting the catalytic active site organization, (Mg^2+^)2 is represented in van der Walls while the protein aminoacids complexing the bimetallic cluster and closest nucleic acid residues are shown in ball and sticks. B) Time series of the evolution of the distance between the phosphate OP atom and the closest Mg^2+^ ion. C) Radial distribution function for the distance between Mg^2+^ and water oxygen atoms.

Indeed, the catalytic activity of bimetallic endonucleases has been shown to proceed through a relatively straightforward mechanism in which one water molecule, complexed to the metallic cluster, attacks the phosphorus, inducing the breaking of one P-O and hence cleaving DNA (Figure 4A). Importantly, nuclease selectively takes place with a 3’→5’ directionality, hence leading to the production of a P-OH and a sugar-O^-^ fragment, respectively. Moreover, the cleavage of the oxygen-phosphorus bond is also favored by the interaction of the phosphate group with the bimetallic cluster, which facilitates the nucleophilic attack of water. The deprotonation and activation of the nucleophilic moiety is also fundamental in favoring the reaction, hence strengthening the importance of proton-acceptor aminoacids in the nearby of the active site.

**Figure 4.**
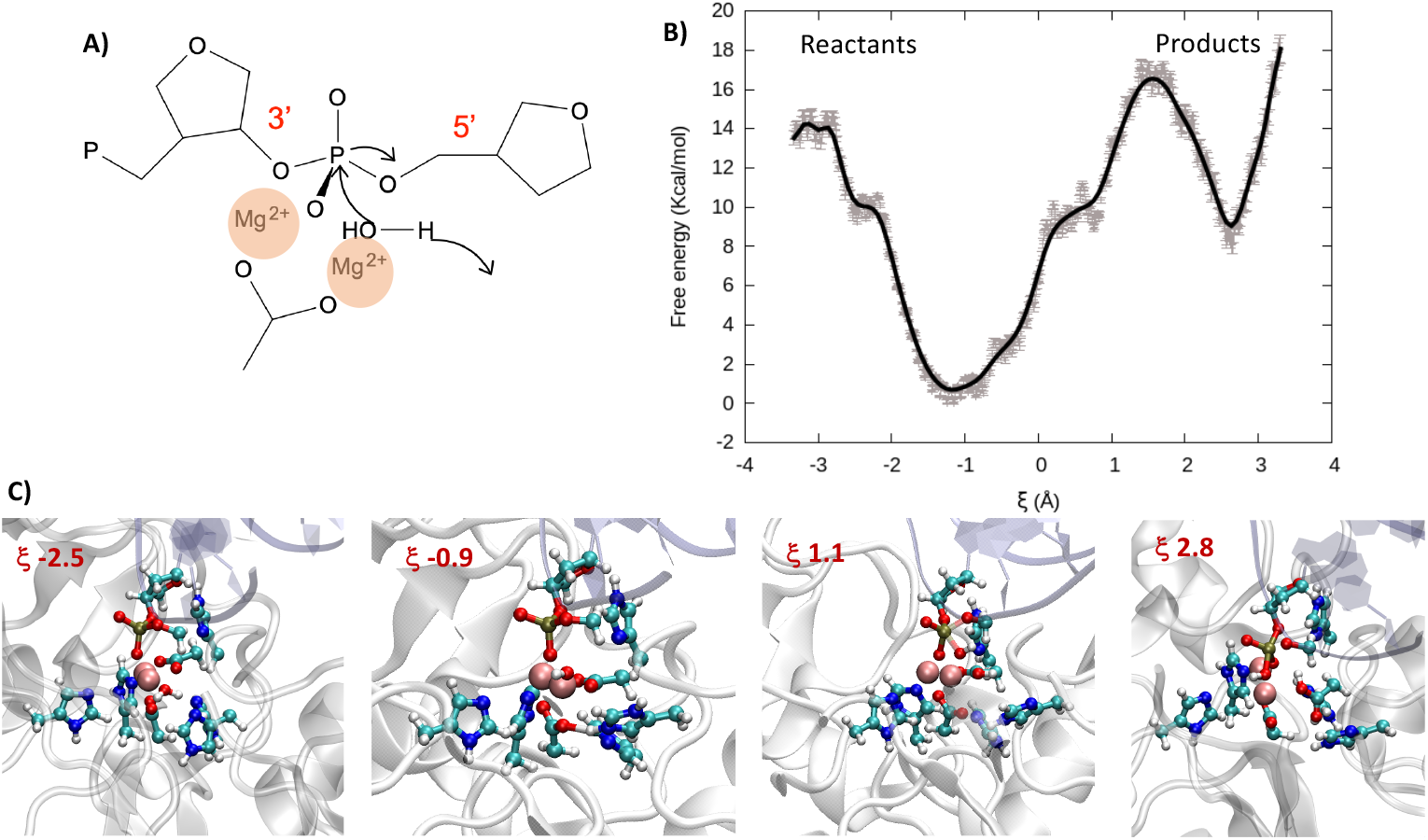
A) Schematic representation of the catalytic attack of a water molecule to the phosphate. B) FEP over the reaction coordinate obtained at DFT level of theory. C) Representative snapshots along the reaction coordinate illustrating the reactant, transition state, and product region. Note that the QM partition is represented in balls and sticks.

In the case of Artemis, the exact nature of the catalytic process is still quite debated, and in particular the capacity of a magnesium cluster of inducing the catalytic reaction and the nature of the deprotonating aminoacids are still controversial [20]. For this reason, and having accessed a reasonable structure for the Artemis/DNA complex we also performed QM/MM enhanced sampling to describe the reaction. Note that, in agreement with the most accepted kinetic model for endonucleases, prior to perform the QM/MM simulations we have forced the DNA phosphate oxygen to reach a distance of only about 3 Å with the (Mg^2+^)_2_ cluster through steered molecular dynamic simulations. For our enhanced sampling simulation, we have considered a reaction coordinate ξ which is defined as the difference of the distance between one hydrogen of the reactive water molecule and the center of mass of the oxygen and nitrogen in the lateral chains of Asp 61 and His 62 and the distance of the oxygen atom of the same water molecule with the reactive phosphorus (See SI for illustration). Hence, while negative values of ξ correspond to the reactant region, positive values indicate the products. As shown in Figure 4B through the FEP and pictorially in the snapshots of Figure 4C the initially Mg-bond water is rapidly deprotonated thanks to the concerted action of Asp 61 and His 62, which participate in the stabilization of the proton. This leads to both an enhanced global stabilization of the system as shown by the shallow minimum at ξ 0.9 and the production of a more nucleophilic species namely the Mg-bond OH^-^ anion. Proceeding further along the reaction coordinate a free energy barrier of about 16 kcal/mol should be overcome to reach the rate-determining transition state (TS) in which the 5’ PO bond is weakened while OH^-^ is approached. As previously observed the FEP basin appears quite shallow and most specifically one can observe a plateau at ξ 0.5 (∼ 10.0 kcal/mol) which is the result of the formation of partially stabilizing interactions between OH^-^ and the residues around the catalytic site. The presence of the plateau can also facilitate overcoming the global free energy barrier and hence increasing the overall catalytic efficiency. Indeed, once the TS is attended the FEP evolves sharply and continuously towards a minimum energy region corresponding to the cleaved backbone. Obviously in this study we are not taking into account neither the protonation of the resulting sugar-O^-^ fragment nor the release of the two cleaved DNA fragments that should globally lower the product free energy due to both enthalpic and entropic factors.

## 3. Discussion and Conclusion

The role of endonucleases in general, and Artemis in particular, is essential in assuring both a proper reparation of DNA lesions and the efficient adaptation and maturation of the immune system. This mostly relies in their efficiency and flexibility in cleave different DNA strands, including hairpins, a flexibility that should find an echo in peculiar structural and reactive features of the enzyme. Furthermore, the role of Artemis and its importance for T and B cells production also make this protein an attractive target for cancer therapy, including radiotherapy sensitization and immunotherapy. Despite all these considerations, the structure of Artemis has been resolved only recently, and important features such as the precise interaction with DNA or the global structure of the active site remained elusive and open to question.

In this contribution, thanks to multiscale molecular modeling and simulation, we provide a first rationalization of the interaction between Artemis and DNA. More precisely, while providing an evidence for the formation of stable complexes with a DNA double strand, we also clearly identify the Lys rich groove as the crucial driving force behind the interaction. We have also evidenced that basic aminoacids are able to penetrate in the groove of the DNA strand, this in turn, provides the possibility of a corkscrew-like movement of the DNA that while maintaining a tight complex may lead different phosphate groups to the catalytic site. On the contrary, and despite a slight bending of the strand, no particular structural deformations of the nucleic acid can be highlighted.

From a biochemical point of view, we have shown, thanks to QM/MM US sampling, that the cleavage reaction may proceed through the assistance of a bimetallic (Mg^2+^)_2_ cluster and the crucial participation of the nearby protein residues. In particular Asp 61 and His 62 are fundamental since not only they are acting as ligand for the metals by they also deprotonate the reactive water molecule, hence producing OH^-^ anion that is further attacking the phosphate group. Interestingly enough, the initial deprotonation of water is accompanied by a considerable stabilization of the system, as shown by the FEP. The excess energy released in this step may, in the following, facilitate overcoming the free energy barrier, and facilitate the reaction. Interestingly, a relatively small plateau prior to the TS is also evident in the FEP, an occurrence that once again may globally facilitate the catalytic cleavage. Our results, pointing to a Mg-based enzyme are also coherent with experimental observation such as the activity of Artemis in magnesium buffers. The suppression of the catalytic activity by mutation of Glu 61 can also be explained due to its fundamental role in the first water deprotonation step.

Our results, are important in providing deeper structural and mechanistic insights on the activity of Artemis and in the future, we plan to extend them in providing suggestions for the rational design of suitable inhibitors. In parallel, we also plan to increase the study of the endo and exonuclease Pantheon, by providing detailed modeling and simulation of the related SNM1 proteins such as SNM1B/Apollo.

## 4. Materials and Methods

The recently reported [20] crystal structure of Artemis catalytic domain was retrieved from the pdb data base (pdb: 6wo0). To obtain an initial structure of the Artemis/DNA complex an ideal B-DNA double strand, having the same sequence as the one used by Karim *et al*. [20] for Artemis crystallography (5’-cacagctgatcgc-3’), was built using the nucleic acid builder (nab) utilities of Amber [23]. The protein and the nucleic acid have been docked using the NPDock webserver utility [24,25]. All the stable poses obtained presented the interaction of the DNA strand in the positively charged groove of the protein and were basically equivalent, hence only the highest scoring pose has been kept for the subsequent molecular simulations.

The docked complex has been solvated in a cubic water box with a buffer of 9 Å and K^+^ cations have been added to ensure electroneutrality. Since only metal atom was present in the catalytic center, the second Mg^2+^ has been manually added and its position inferred by superposing the crystal structure of Artemis with the one of Apollo exonuclease (pdb: 5aho) for which a totally resolved active site is provided [17]. The protonation state of the amphipathic residues has been determined according to their pKa, estimated with propka [26], with the exception of the His and Cys present in the catalytic site and in a Zn-finger region whose state has been assigned to maximize the interactions with the metals. The protein and the nucleic acid have been modelled with amberf99sb force field [27], including the bsc1 corrections for DNA [28,29], while water is modelled using TIP3P [30]. The aminoacids composing the Zn-finger domain have been described using a recently developed amber-based non-bonded potential [31]. To assure the stability of the bimetallic catalytic site additional constraints to the Mg-Mg distance as well as to the distance of Mg^2+^ with the nitrogen atom of the first-shell His residues have been applied. Two independent replicas have been constructed and MD simulation has been performed using the NAMD code [32,33] in the constant temperature (300 K) and pressure (1 atm) thermodynamic ensemble (NPT) enforced using the Langevin thermostat [34] and barometer [35]. After 5000 minimization steps, the system has been equilibrated for 9 ns during which constraints on the heavy atoms have been progressively released, and finally a 2.0 μs production run for the first replica and 2.5 μs for the second replica have been performed. The use of hydrogen mass reparation (HMR) strategy [36] in combination with the Rattle and Shake procedure [37] has allowed to use of a 4 fs timestep. Particle mesh Ewald (PME) [38] has been used to treat long-range interactions with a cut-off of 9 Å.

For the determination of the reaction mechanism electrostatic embedding QM/MM strategy [39] has been followed, after having preliminary decreased the phosphate-Mg distance by steered molecular dynamics. The chosen QM partition involved: the (Mg^2+^)_2_ cluster; the reactive water molecule; the lateral chains of His57, His59, His62, His139, His343, Asp61, and Asp160; the 3’ phosphate and sugar moiety; and the 5’ OCH_2_ group. Dangling bonds have been saturated with the link-atom approach, hence adding hydrogens, leading to a total of 100 QM atoms. After equilibration the US was performed partitioning the reaction coordinates in 67 windows spaced by 0.1 Å and spanning the -3.3 / 3.3 Å domain, each window has been preliminary equilibrated for 500 fs. QM calculation has been performed at density functional theory (DFT) level using the ωB97X functional [40] and the 6-31G basis set, a time step of 1 fs has been consistently used, while Rattle and Shake has been removed for the QM partition. Note that dispersion has not been taken into account explicitly in DFT to avoid double counting due to the inclusion of van der Waals terms in the QM/MM Hamiltonian. QM/MM calculations have been performed using the Amber/Terachem interface [41–43]. The FEP has been reconstructed using the weighted histogram analysis method (WHAM) algorithm [44,45], the overlapping between the windows has been checked and can be appreciated in SI. Trajectories issued from both classical and QM/MM simulations have been visualized and analyzed using VMD [46], and the structural parameters of DNA are obtained using Curves+ [47].

## Supporting information

Video of the MD trajectory

## Supplementary Materials

Figure S1 RMSD for the second replica, Figure S2 representative snapshots of the trajectory, Figure S3 RMSD for the isolated protein, Figure S4 DNA structural parameters obtained with Curves+, Figure S5 representation of the QM partition, Figure S6 representation of the reaction coordinate, Figure S7 overlap between the umbrella sampling windows. The following are available online at www.mdpi.com/xxx/s1. Video S1 Trajectory of the MD simulation for the Artemis/DNA simulation: artemis_video.mpg.

## Acknowledgments

Calculations have been performed on the local LPCT computing cluster and on the regional Explor computing center under the project “Dancing under the light”.

