## Supplementary Information for "Staring at the naked Goddess. Unraveling structure and reactivity of Artemis endonuclease interacting with a DNA double strand"

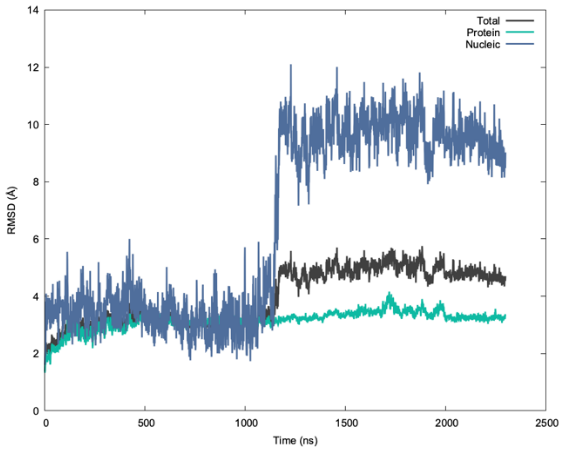


**Figure S1.** RMSD calculated on the second independent replica for the total complex, the protein and the DNA, respectively.

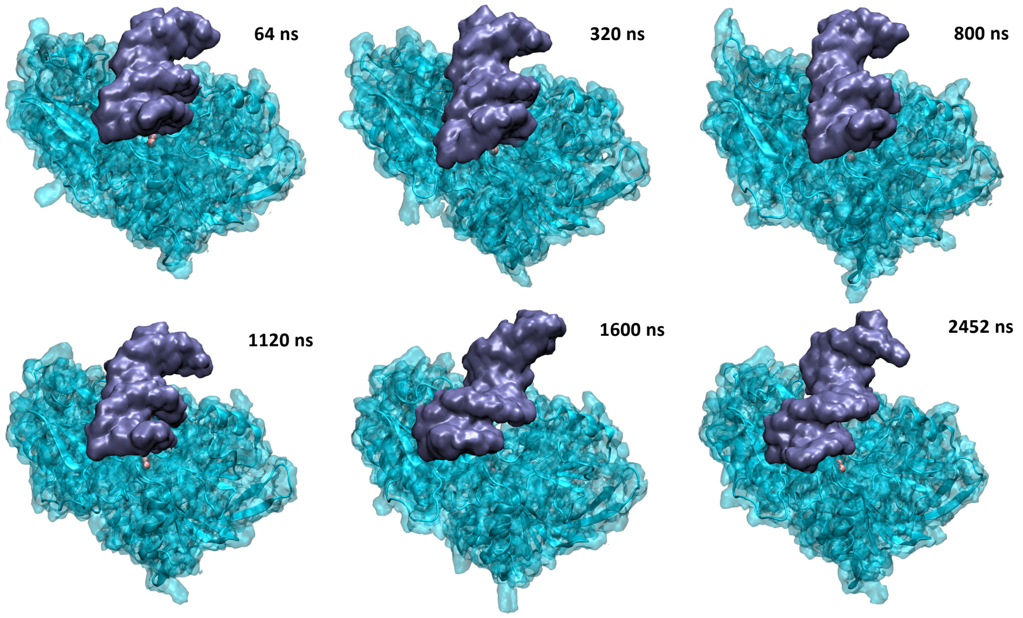


**Figure S2**. Representative snapshots extracted from the trajectory of the second replica.

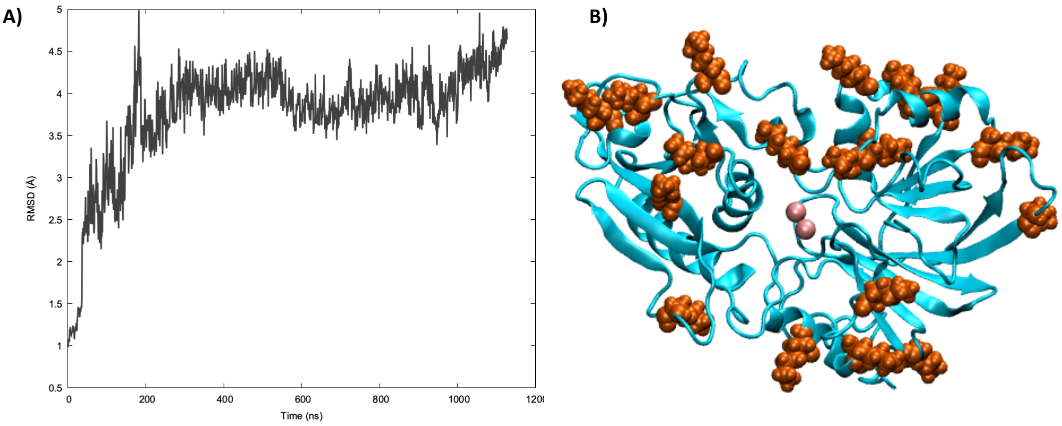


**Figure S3.** A) RMSD calculated for the trajectory of the isolated protein and B) a representative snapshot of the MD simulation showing the main secondary structure feature of Artemis. Note that the positively charged Lys are highlighted in van der Waals representation and in orange, while the $(Mg^{2+})_2$ cluster is shown in pink.

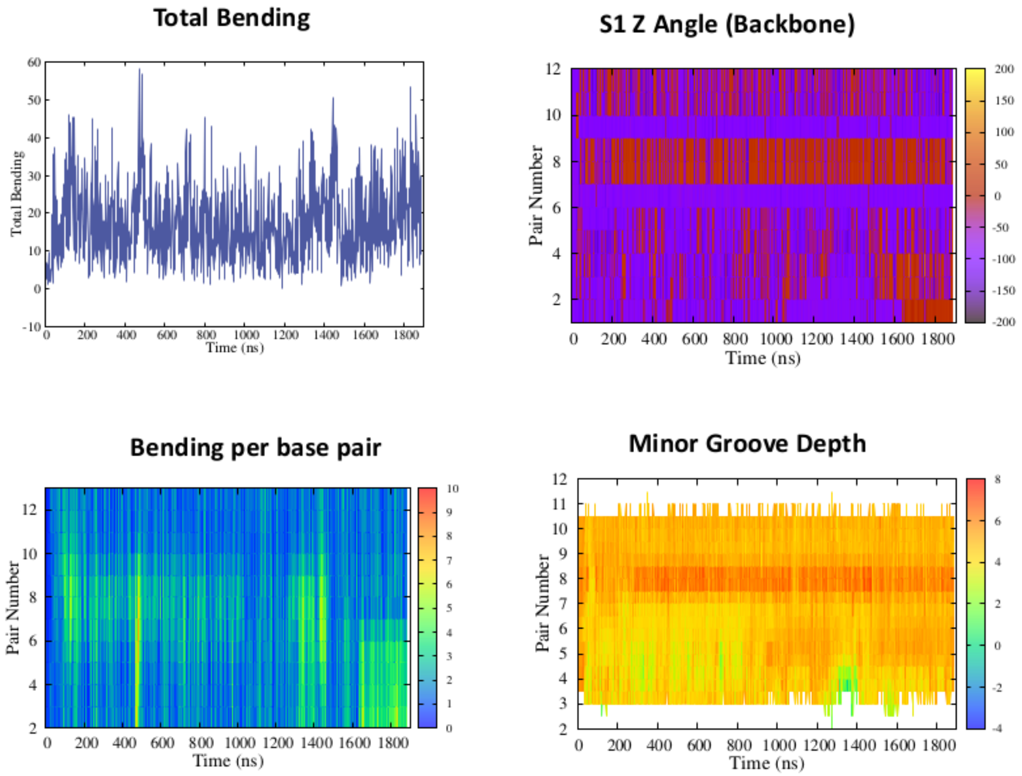


**Figure S4.** Selected Curves+ parameters extracted from the trajectory of the first replica. Note the slight perturbation of the parameters in correspondence of the base pair 8, i.e. the phosphate exposed to the catalytic site.

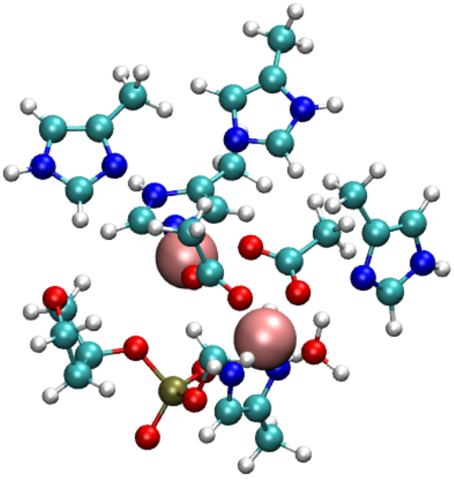

**Figure S5.** Representation in ball and stick and van der Waals of the QM partition, including the hydrogen link atoms.

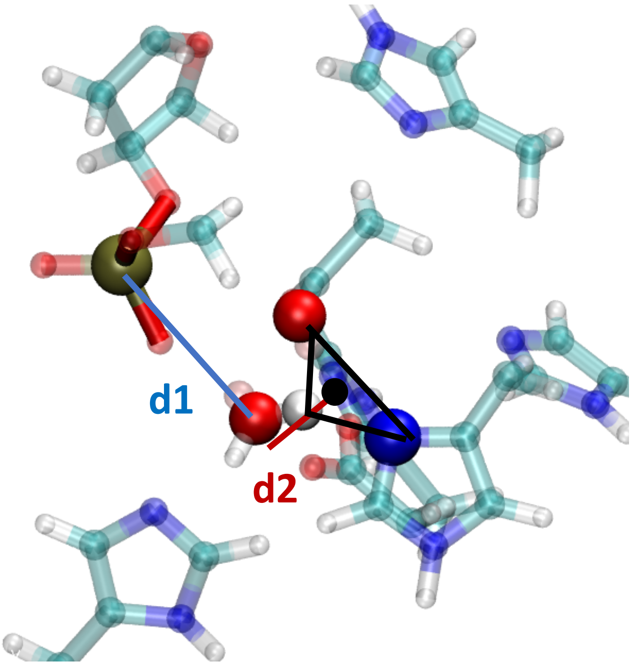


**Figure S6.** Representation of the reaction coordinate $\xi$ = d2 – d1. Note that d2 is calculated as the distance between the water oxygen and the center of mass of the highlighted water hydrogen, nitrogen of Hys, and oxygen of Asp. Negative values correspond to the reactants while positive values indicate the product region.

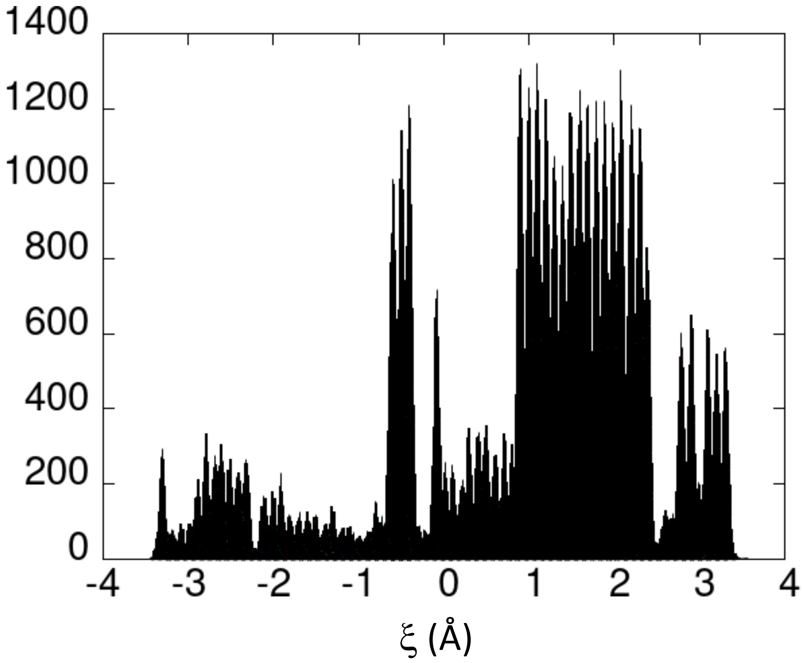


**Figure S7.** Distribution of the reaction coordinate x among the different windows.
